# CyanOperon: an operon building expansion for the CyanoGate MoClo toolkit

**DOI:** 10.64898/2026.02.24.707249

**Authors:** Michael J. Astbury, Alejandra A. Schiavon Osorio, Angelo J. Victoria, Alistair J. McCormick

## Abstract

Operons are gene clusters controlled by a single promoter that enable coordinated translation from a single messenger RNA. Here we describe an expansion of the CyanoGate MoClo toolkit to assemble synthetic operons. The versatile CyanOperon system includes two Level 0 acceptor vectors for building interchangeable promoter-ribosome bind site (RBS) combinations and 15 Level 1 acceptor vectors for the hierarchical assembly and expression of up to six genes within a single operon. The system also allows for operon assembly into a self-replicating vector or for chromosomal integration by homologous recombination. To showcase CyanOperon, we assembled the violacein biosynthesis pathway as an operon and demonstrated violacein production in *Escherichia coli*. We then constructed a 20-part RBS library to examine how spacer length between the Shine–Dalgarno sequence and start codon affects translation in *E. coli* and the model cyanobacterium *Synechocystis* sp. PCC 6803. Lastly, we compared the expression of up to three operonic fluorescent markers following chromosomal integration or from a self-replicating vector in *E. coli* and *Synechocystis* sp. PCC 6803. The CyanOperon system is publicly available and can be readily integrated with other MoClo systems to accelerate the development of standardized operon assemblies.

**GRAPHICAL ABSTRACT:** 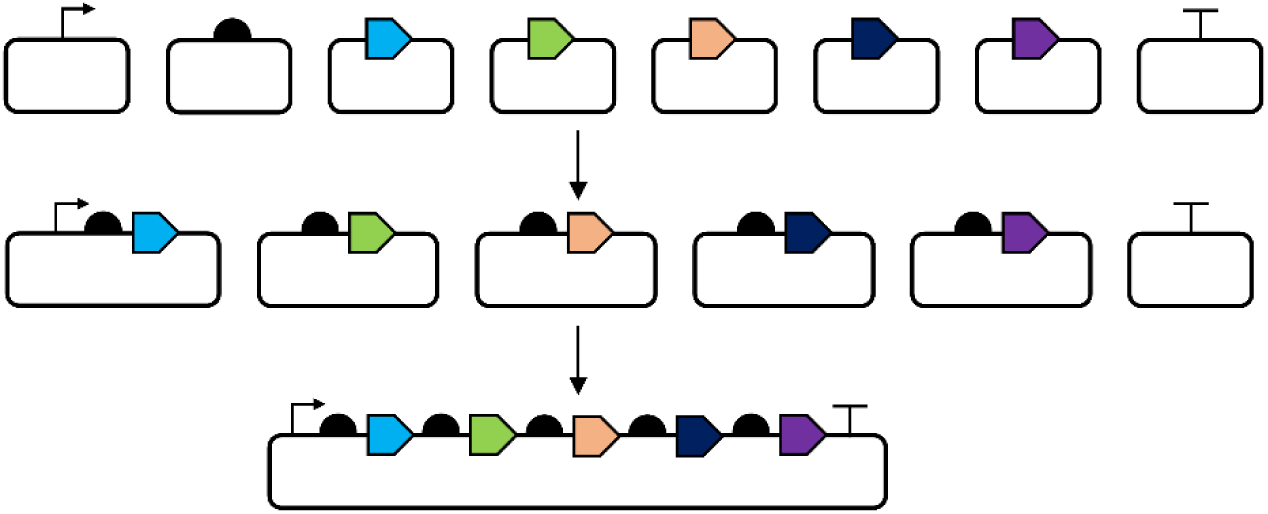

## Introduction

Engineering biology relies on the availability of modular DNA parts (e.g., promoters, coding sequences (CDSs) and terminators) to assemble molecular devices (e.g., functional transcriptional units), which can then be combined into larger genetic modules (e.g., biosynthetic pathways). To date, several modular DNA assembly methods have been developed using a Type IIS restriction endonuclease-mediated ‘MoClo’ approach (Engler et al., 2014; Taylor et al., 2021; Valenzuela-Ortega and French, 2021; Blázquez et al, 2023; Inckemann et al. 2025). Most of these rely on a common syntax of fusion sites to standardize the assembly of modular parts (Patron et al., 2015).

In recent years, much progress has been made towards developing synthetic biology tools for cyanobacteria (Markley et al., 2015; Englund et al., 2016; Sun et al., 2018; Wang et al., 2018; Vasudevan et al., 2019; Opel et al., 2022; Victoria et al., 2024). Such approaches have enabled metabolic engineering strategies for the production of alcohols (Gao et al., 2012; Oliver et al., 2014; Namakoshi et al., 2016), sugars (Lin et al., 2020; Domínguez-Lobo e al., 2025), terpenoids (Lin et al., 2017; Englund et al., 2018; Rodrigues et al., 2021; Rohr and Rudroff, 2025) and fatty acids (Wlodarczyk et al., 2020). Several operon-based metabolic engineering strategies using polycistronic constructs have been demonstrated in cyanobacteria (e.g., Wlodarczyka et al., 2016; Brey et al., 2017; Rodrigues et al., 2021), but a standardized hierarchical system for the design and assembly of synthetic operons is still lacking in the cyanobacterial community (Till et al., 2020).

In prokaryotes, clustering of functionally related genes into operons allows for simultaneous and coordinated gene expression and translation (Koonin, 2009). Up to 50% of genes are predicted to be transcribed as part of an operon in a typical prokaryotic genome (Memon et al., 2013). Operons typically comprise multiple coding sequences, each with its own 5′ untranslated region containing a ribosome binding site (RBS; 15–20 nucleotides (nt)), transcribed as a single polycistronic mRNA. Differential expression of operon-encoded proteins can be achieved through the regulation of translation efficiency by intercistronic sites (e.g., RBSs), which together contribute to the modulation of individual protein expression levels (Zelcbuch et al., 2013). Translation initiation in prokaryotes typically proceeds by binding of the 30S ribosomal subunit to an RBS on the mRNA. This is generally dependent on the Shine-Dalgarno (SD) sequence within the RBS, which base pairs with the anti-SD sequence in 16S ribosomal RNA (rRNA) component of the 30S ribosomal subunit to guide selection of the appropriate downstream start codon (Shine and Dalgarno, 1974). Key determinants of translation initiation rate include (i) SD/anti-SD pairing strength, (ii) SD–start codon spacing, and (iii) mRNA secondary structures affecting RBS accessibility (Studer and Joseph, 2006; Allert et al., 2010; Quax et al., 2013).

Translation initiation in prokaryotes is a key rate-limiting step in regulating protein expression Levels (Nakagawa et al., 2010; Wei and Xia, 2019). Thus, engineering RBS sequences offers the ability to modulate metabolism with just a few nucleotide changes. In *Escherichia coli* (*E. coli* hereafter), RBS sequences have been extensively studied to determine the molecular interactions that affect protein translation (Marshall and Noireaux, 2019). The use of *in silico* tools for rational design of RBS sequences is now a common practice in *E. coli* synthetic biology (Salis et al., 2009). In cyanobacteria, several studies have characterised native, heterologous, and synthetic RBS sequences to investigate their impact on protein translation (Heidorn et al., 2011; Markley et al., 2015; Englund et al., 2016; Lui and Pakrasi, 2018; Thiel et al., 2018; Wang et al., 2018; Wei and Xia, 2019). However, the predictive power of *in silico* tools does not yet extend well to cyanobacteria, as the observed *in vivo* activities of *in silico* designed RBSs have correlated poorly with the predicted activity (Oliver et al., 2014; Markley et al., 2015; Wang et al., 2018; Taylor et al., 2021). More recently, large-scale library screening approaches have been used to identify RBS-coding sequence configurations for optimal gene expression (Jodlbauer et al., 2024).

Here we have designed, developed and tested an expansion of the CyanoGate MoClo toolkit for the straightforward hierarchical assembly of promoters, RBSs and gene coding sequences into synthetic operons for expression in bacterial or cyanobacterial hosts (Engler at al., 2014; Vasudevan et al., 2019). Below we describe the design of the CyanOperon system and highlight three proof-of-concept experiments conceived to showcase its utility.

## Results and Discussion

### Design and construction of new acceptor vectors and parts for CyanOperon

The CyanOperon system builds on the CyanoGate and Golden Gate MoClo Plant (Engler et al., 2014; Vasudevan et al., 2019) toolkits, which can be acquired from Addgene (see Vasudevan et al., 2019 for details). We have retained the syntax of Level 0 parts (i.e., all existing Level 0 parts are compatible) and introduced two new Level 0 acceptor vectors to allow for separate assembly of i) core promoter sequences truncated to the transcription start site (‘TSS promoter’ parts, with MoClo overhangs GGAG-TACT) and ii) ribosome binding sites and spacers prior to the start codon (‘RBS parts’, with MoClo overhangs TACT-AATG) (**Table 1**, **Fig. 1A**).

**Figure 1.**
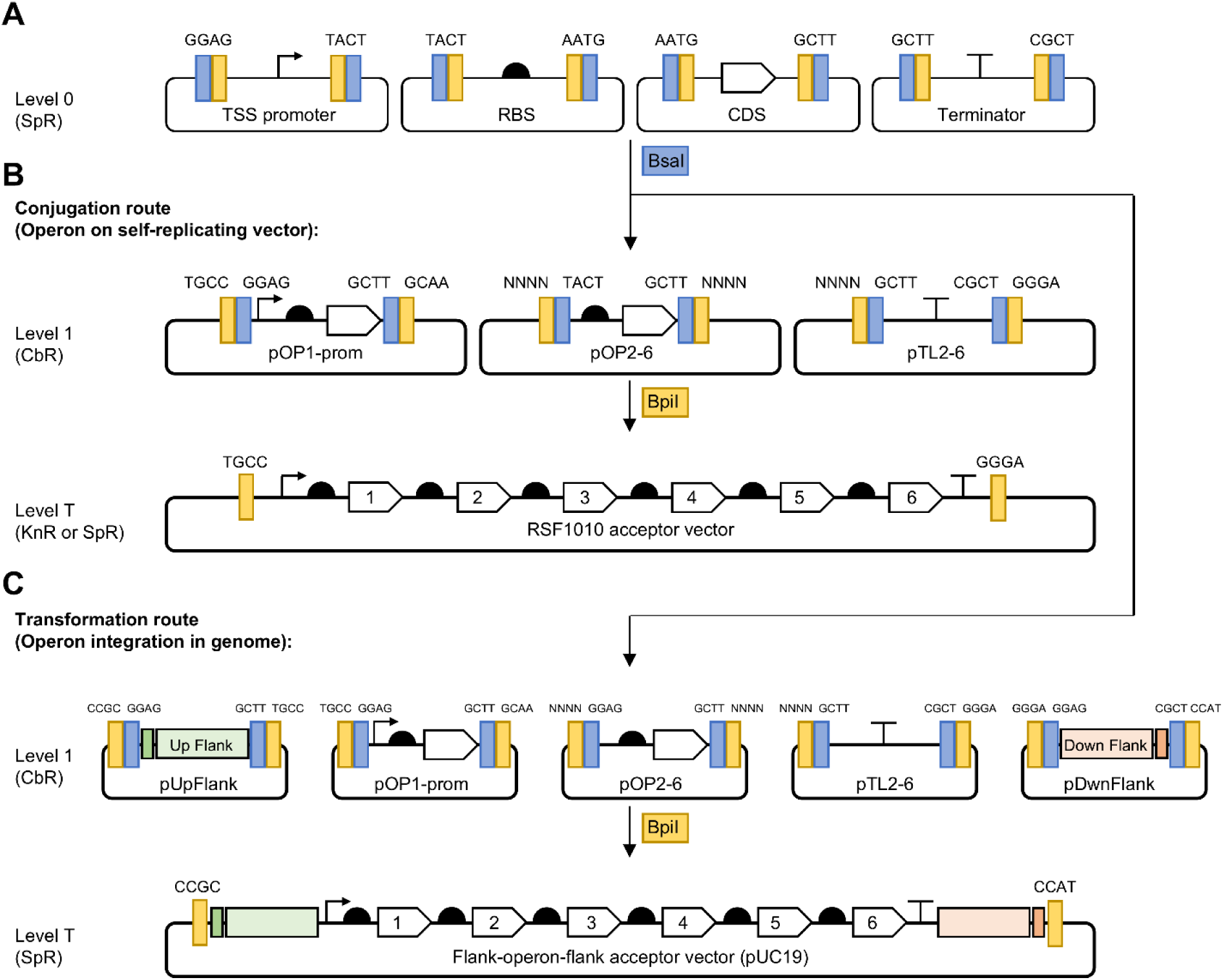
CyanOperon extension of the CyanoGate platform for generating operons. (**A**) Syntax of Level 0 parts for the TSS promoter, RBS, coding sequence (CDS) and terminator. All Level 0 acceptor vectors have internal BpiI (yellow) restriction sites to allow for uptake of Level 0 parts and external BsaI (blue) restriction sites for assembly into Level 1 acceptors (**Table 1**). (**B**) New Level 1 acceptor vectors for operon expression. The acceptor vector pOP1-prom was designed to contain internal overhangs GGAG-GCTT to accommodate a TSS promoter, RBS and CDS (i.e., the first gene in the operon). Acceptor vectors pOP2 to pOP6 contain internal overhangs TACT-GCTT to accommodate an RBS and a CDS (i.e., 2 to 6 subsequent genes). The pTL2 to pTL6 acceptor vectors can accept a terminator part (GCTT-CGCT) and act as End Linkers for pOP2 to pOP6, respectively, to allow for assembly into a Level T acceptor. ‘NNNN’ represents the unique BpiI overhang for each acceptor vector (see **Table 1**, **Fig. S1**). Shown at the bottom of (B) is an operon assembly in a self-replicating acceptor vector (e.g., the RSF1010-derived vector pPMQAK1-T) for operon expression in conjugation-compatible cyanobacterial species. (**C**) Two new acceptor vectors facilitate genomic integration of operons by introducing upstream (pUpFlank) and downstream (pDwnFlank) flank sequences for homologous recombination. pUpFlank and pDwnFlank are designed to accept a Level 0 flank and linker part from the CyanoGate system. A pUC19-based Level T Flank-operon-flank acceptor vector allows for assembly and transformation.

**Table 1.** List of available plasmid vectors in the CyanOperon system. Vectors pCA0.324 and pCA0.325 are acceptor vectors for promoters truncated to the transcription start site (TSS promoter) and ribosome binding site (RBS) (**Fig. 1A**). Note the different overhangs for the TSS acceptor (GGAG-TACT) compared to the TSS acceptor in the CyanoGate system (GGAG-TAGC) (Vasudevan et al., 2019). The new overhang now complies with the 3’ syntax standard for the CORE promoter region (Patron et al., 2015). pCA1.261-pCA1.273 facilitate operon assembly for expression of up to six gene coding sequences (**Fig. 1B**). Vectors pCA1.552 (pOP2-prom) can be used in place of pCA1.552 (pOP1-prom) if an alternative Level 1 vector is being assembled at position 1 (e.g. a separate gene expression cassette). Level 1 parts can be assembled together into a compatible Level T acceptor vector, such as the CyanoGate RSF1010-derived pPMQAK1-T (pCAT.000) vector (Vasudevan et al., 2019). Level 1 acceptor vectors pCA1.620 and pCA1.644 facilitate chromosomal integration of the operon by introducing upstream (pUpFlank) and downstream (pDwnFlank) flank sequences for homologous recombination (**Fig. 1C**). Level T acceptor pCAT.703 allows for assembly of operon vectors, with pUpFlank and pDwnFlank vectors for chromosomal integration. All acceptor vectors are compatible with blue/white screening and contain a spectinomycin resistance cassette for Level 0 and T, and a carbenicillin resistance cassette for Level 1. Plasmid maps are in **Materials S1** and available on Addgene.

| Vector Name | Addgene ID | Plasmid name | Description | Level | Overhangs (Bsal) | Overhangs (Bpil) |
| --- | --- | --- | --- | --- | --- | --- |
| **Acceptor vectors** |  |  |  |  |  |  |
| pCA0.324 | 200621 | TSS acceptor | for TSS promoter parts | 0 | GGAG - TACT |  |
| pCA0.325 | 200622 | RBS acceptor | For RBS parts | 0 | TACT - AATG |  |
| pCA1.261 | 200607 | pOP1-prom | Operon acceptor position 1 | 1 | GGAG - GCTT | TGCC - GCAA |
| pCA1.262 | 200608 | pOP2 | Operon acceptor position 2 | 1 | TACT - GCTT | GCAA - ACTA |
| pCA1.552 | 220334 | pOP2-prom | Operon acceptor position 2<br>(if L1P1 occupied) | 1 | GGAG - GCTT | GCAA - ACTA |
| pCA1.263 | 200609 | pOP3 | Operon acceptor position 3 | 1 | TACT - GCTT | ACTA - TTAC |
| pCA1.264 | 200610 | pOP4 | Operon acceptor position 4 | 1 | TACT - GCTT | TTAC - CAGA |
| pCA1.265 | 200611 | pOP5 | Operon acceptor position 5 | 1 | TACT - GCTT | CAGA - TGTG |
| pCA1.266 | 200612 | pOP6 | Operon acceptor position 6 | 1 | TACT - GCTT | TGTG - GAGC |
| pCA1.268 | 200614 | pTL1 | Terminator end-linker position 1 | 1 | GCTT - CGCT | GCAA - GGGA |
| pCA1.269 | 200615 | pTL2 | Terminator end-linker position 2 | 1 | GCTT - CGCT | ACTA - GGGA |
| pCA1.270 | 200616 | pTL3 | Terminator end-linker position 3 | 1 | GCTT - CGCT | TTAC - GGGA |
| pCA1.271 | 200617 | pTL4 | Terminator end-linker position 4 | 1 | GCTT - CGCT | CAGA - GGGA |
| pCA1.272 | 200618 | pTL5 | Terminator end-linker position 5 | 1 | GCTT - CGCT | TGTG - GGGA |
| pCA1.273 | 200619 | pTL6 | Terminator end-linker position 6 | 1 | GCTT - CGCT | GAGC - GGGA |
| **Acceptor vectors for integration** |  |  |  |  |  |  |
| pCA1.620 | 240219 | pUpFlank | Up Flank acceptor | 1 | GGAG- GCTT | CCGC-TGCC |
| pCA1.644 | 240218 | pDwnFlank | Down Flank acceptor | 1 | GGAG-CGCT | GGGA-CCAT |
| pCAT.703 | 240217 | pUC19-T | Flank-operon-flank acceptor | T | - | CCGC-CCAT |
| **Level 0 parts** |  |  |  |  |  |  |
| pC0.335 | 200623 | P<sub>V02</sub> TSS |  | 0 | GGAG - TACT |  |
| pC0.336 | 200624 | P<sub>J23100</sub> TSS |  | 0 | GGAG - TACT |  |
| pC0.337 | 200625 | P<sub>J23101</sub> TSS |  | 0 | GGAG - TACT |  |
| pC0.338 | 200626 | P<sub>Trc10</sub> TSS |  | 0 | GGAG - TACT |  |
| pC0.340 | 200628 | RBS* |  | 0 | TACT - AATG |  |
| pC0.341 | 200629 | RBS B0034 |  | 0 | TACT - AATG |  |

Level 1 acceptor vectors typically require the inclusion of a Level 0 terminator part (GCTT-CGCT) for assembly into a transcriptional unit (Engler et al., 2014). To allow for the assembly of operonic transcriptional units that are driven by only an RBS and lack a terminator, we designed a suite of new Level 1 acceptor vectors (**Table 1**, **Fig. 1B**). The first transcriptional unit is assembled in the Level 1 acceptor vector pOP1-prom with overhangs GGAG-GCTT, so that a promoter and gene coding sequences can be assembled without a terminator. The subsequent five Level 1 acceptor vectors pOP2-6 contain overhangs TACT-GCTT for assembly of an RBS and a gene coding sequence. The BpiI and BsaI restriction sites for the pOP1-6 acceptor vectors were placed together so that the 4 bp overhangs were adjacent each other (**Fig. S1**). As such, the sequence length between the upstream gene stop codon and downstream RBS was reduced to 12 bp for each operonic transcriptional unit. We generated a matching suite of Level 1 ‘terminator end-linker’ vectors (pTL2-6) that can each be assembled with a Level 0 terminator part of choice to complete the operon in Level T. For example, assembled pOP1-prom, pOP2, pOP3 and Terminator end-linker position 3 (pTL3) vectors can be combined together into a Level T acceptor vector to generate an operon consisting of three gene coding sequences. Thus, the choice of the core promoter, the RBS driving each gene coding sequences and the terminator part within the operon can be readily exchanged using standard MoClo swapping and re-assembling strategies.

The new suite of Level 1 acceptor and terminator end-linker vectors are compatible with and can assemble into existing CyanoGate Level T acceptors vectors (e.g., the RSF1010-derived pPMQAK1-T vector) (**Fig. 1B**). To allow for targeted genomic integration by homologous recombination we generated two additional Level 1 acceptors vectors (pUpFlank and pDwnFlank) that enable assembly of Level 0 Up Flank and Down Flank parts, respectively (**Table 1**, **Fig. 1C**) (Vasudevan et al., 2019; Victoria et al., 2024). pUpFlank and pDwnFlank vectors can then be assembled with pOP and pTL vectors into the Level T vector pUC19-T, which subsequently can be used for genomic integration.

### Assembly of the violacein pathway using the CyanOperon system

To validate that the CyanOperon system can generate a functional multi-gene operon, we assembled the VioABCDE operon and tested for violacein production in *E. coli*. Violacein is a deep-purple bis-indole pigment primarily produced by *Chromobacterium* species (Pemberton et al., 1991), and its metabolic production pathway is commonly used in *E. coli* as a model for metabolic engineering (Rodrigues et al., 2014; Jones et al., 2015; Immanuel et al., 2018; Mehta et al., 2025). The pathway has previously been demonstrated to function in yeast following MoClo-based assembly of separate gene expression cassettes for the five pathway components (i.e., VioA-VioE) (Tong et al., 2021), but not yet for an operon-based MoClo system.

We sourced the five genes of the violacein operon and cloned these separately into the Level 0 the CDS1 acceptor vector (pICH41308) (**Fig. 2A**) (Engler et al., 2014). We then constructed Level 0 parts for six TSS promoters of different putative strengths (P_J23100 TSS_, P_J23101 TSS_, P_J23105 TSS_, P_J23107 TSS_, P_J23110 TSS_ and P_V02 TSS_) (Vasudevan et al., 2019) by annealing synthesised oligonucleotides and assembling these into the TSS acceptor vector (**Table 1**; **Table S1**). For Level 1 assemblies, each TSS promoter part was assembled with RBS part B0034 (pC0.341), derived from the broad-range BioBricks BBa_B0034 RBS, and the Level 0 VioA part into the pOP1-prom acceptor vector. VioB, VioE, VioD and VioC were each assembled with RBS B0034 into the pOP2 – pOP5 acceptor vectors, respectively. The T_rrnB_ double terminator part (pC0.082; Vasudevan et al., 2019) was assembled into pTL5. Finally, the five genes were assembled into the pPMQAK1-T acceptor vector to generate six different Level T vectors where the VioABCDE operon was driven by different promoters (**Materials S2**).

**Figure 2.**
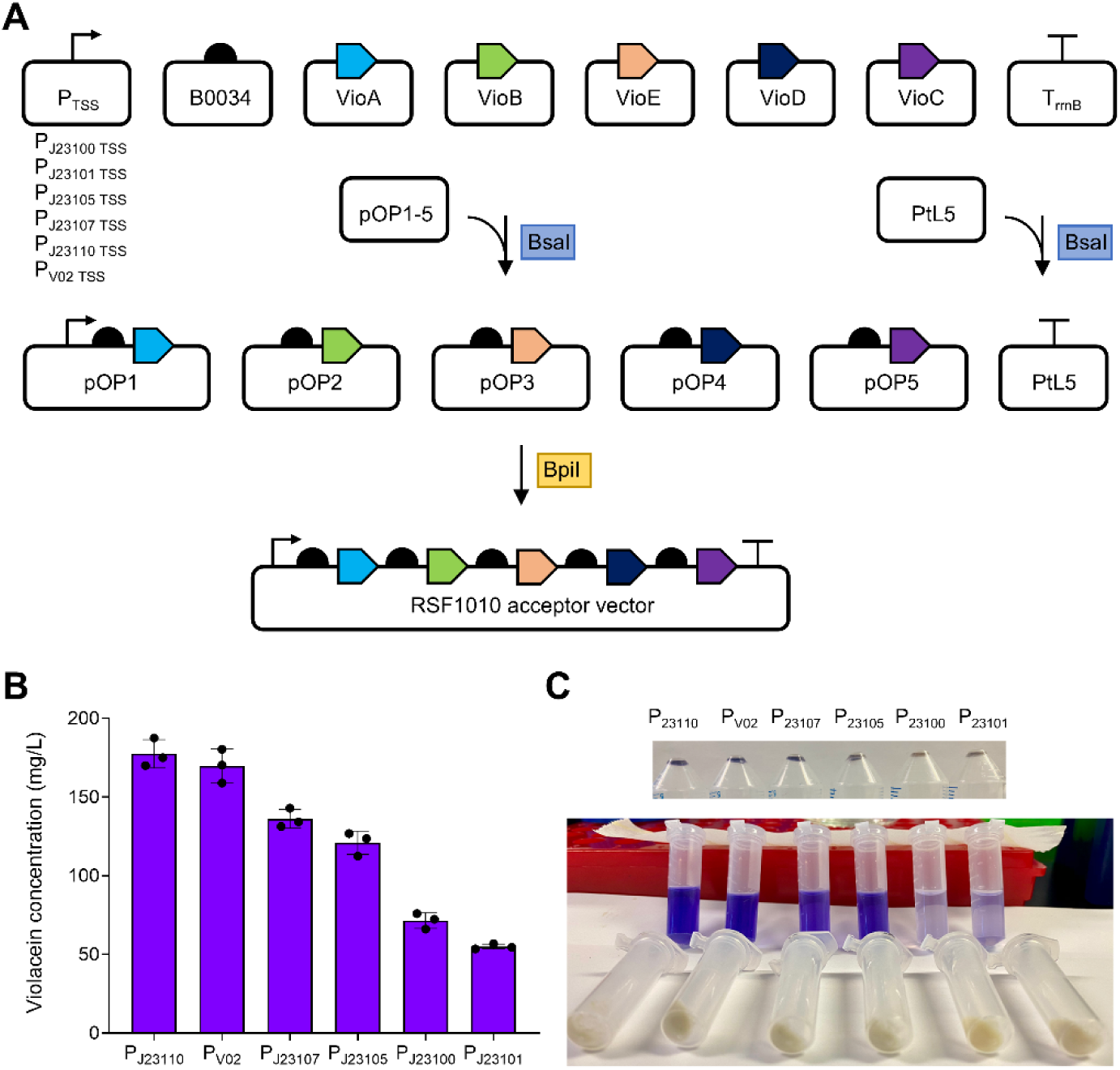
Assembly and validation of the violacein operon in *E. coli*. Violacein is synthesised by the sequential activities of the five enzymes produced by the VioABCDE operon (Tong et al., 2021). The amino acid substrate L-tryptophan is converted into indole-3-pyruvic acid imine (IPA imine) by VioA, which is then dimerised by action of VioB. The IPA imine dimer is converted into pro-todeoxyviolaceinic acid by VioE. VioD then catalyses the conversion of protodeoxyviolaceinic acid into protoviolaceinic acid. The final deep purple product, violacein, results from conversion of protoviolaceinic acid via the action of VioC. (**A**) The five components of the violacein pathway were assembled using pOP1-prom and pOP2-5, and pTL5 carrying the T_rrnB_ double terminator part (pC0.082) (Vasudevan et al., 2019). (**B**) Violacein content follow expression with six different TSS promoter parts. (**C**) Examples of violacein content before (top) and after (bottom) extraction.

We observed that operons driven by lower strength promoters generally produced higher violacein yield than those driven by stronger promoters in *E. coli*. The low strength promoter P_J23110_ produced the highest violacein yields (178 mg/L), while the P_J23100_ and P_J23101_, producing 71 mg/L and 55 mg/L, respectively (**Fig. 2B**). These findings are consistent with previous work in *E. coli*, where reduced strength promoters correlated with higher violacein yields (Jones et al., 2015), while stronger promoters can result in feedback inhibition, possibly on aromatic compound production (Li et al., 2020). These proof-of-concept results demonstrate that CyanOperon could be used to rapidly optimize promoter and RBS strengths using MoClo-based combinatorial optimization strategies (Jescheck et al. 2017; Jodlbauer et al., 2024).

We subsequently conjugated the Level T vectors into the model cyanobacterium *Synechocystis* sp. PCC 6803 (*Synechocystis* hereafter) but failed to detect violacein production. Recent work using the native or a refactored violacein biosynthetic gene cluster from the marine γ-proteobacterium *Pseudoalteromonas luteoviolacea* did report low yields of violacein when expressed in *Synechocystis* (1-6 mg/L) (Dhakal et al. 2025), suggesting that our operon could be adjusted to improve production. The low reported yields may also be due to limited availability of the aromatic amino acid substrate tryptophan to drive production (Pemberton et al., 1991). However, tryptophan overproduction strains have been produced in *Synechocystis* (Deshpande et al. 2020). Recent efforts to produce betaxanthin in *Synechocystis* also failed to detect the product until the shikimate pathway was engineered to over accumulate aromatic amino acids, including tryptophan (Hanamghar et al., 2025). Together, these studies indicate potential future strategies for further increasing violacein production in cyanobacteria.

### Investigating the impact of the RBS spacer region on translation

We next used CyanOperon to examine how spacer length between the Shine–Dalgarno (SD) sequence and start codon affects translation efficiency, using eYFP as a reporter. Using the RBS part B0034 as a template, which contains the core SD sequence AGGAGA, a subset of 16 variants was generated in which the spacer length between the SD sequence and the start codon varied from 0 to 14 nt (**Fig. 3A**). Additional RBSs tested included the BioBrick standards B0032 and B0033, the RBS sequence for the *Synechocystis cpc* operon (RBS cpc, taken as the sequence 30 nt upstream of the *cpcB* start codon) and RBS* (Heidorn et al., 2011; Zhou et al., 2014). Each RBS was assembled with the P_V02 TSS_ promoter (pC0.335), derived from a previously characterised P_J23119_ variant, 23119MH_V02, which drives robust expression in *Synechocystis* (Vasudevan et al., 2019), and with eYFP to generate the expression cassette format P_V02 TSS_–RBSx–eYFP–T_rrnB_ (**Fig. S2A**). The Level 1 cassettes were then cloned into the RSF1010-derived pPMQAK1-T vector for characterisation in *E. coli* and *Synechocystis*.

**Figure 3.**
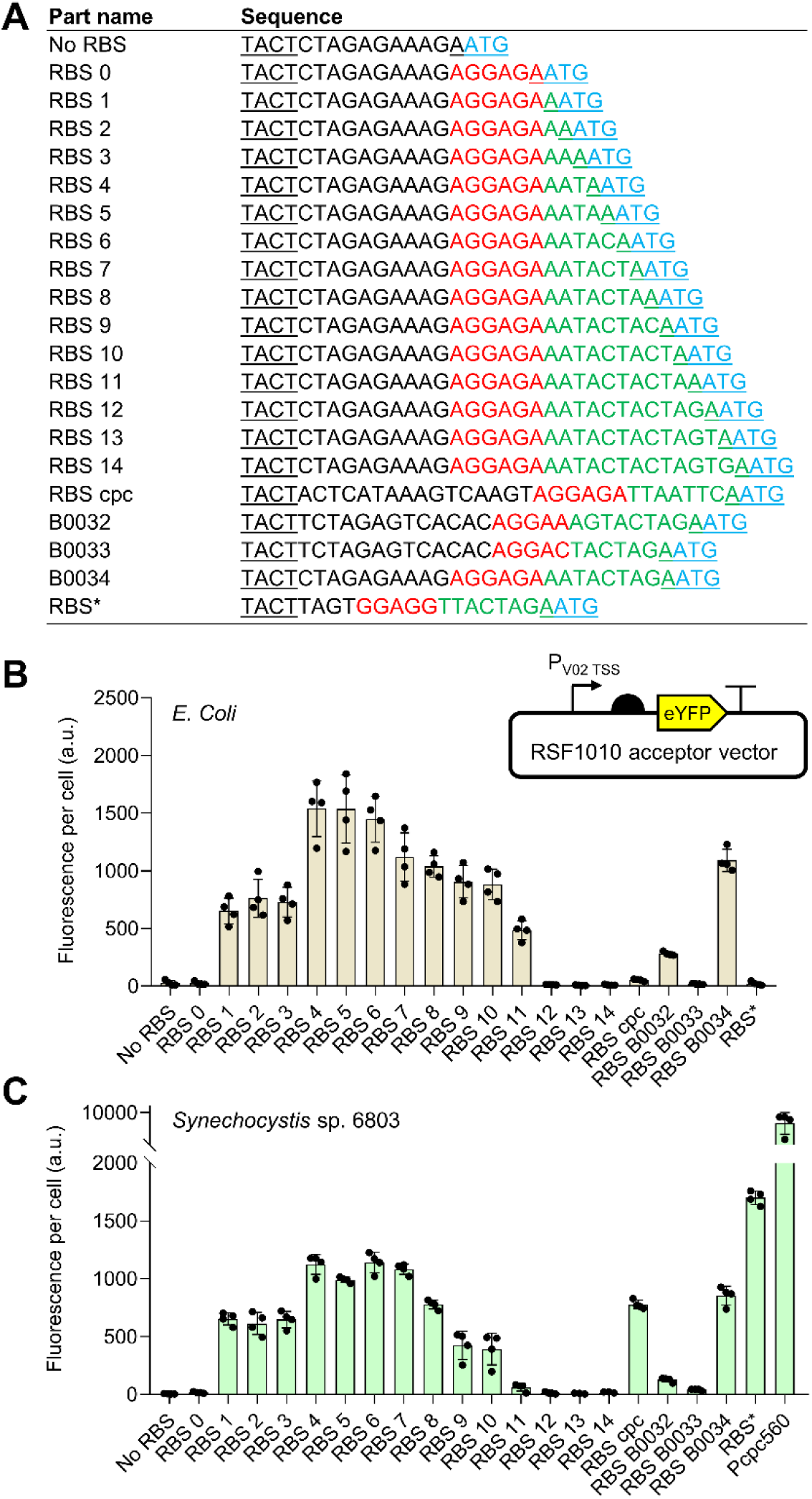
Ribosome bind site parts and their performance in *E. coli* and *Synechocystis* sp. PCC 6803. (**A**) Each RBS sequence is annotated with the adjacent MoClo overhangs (underlined), the SD sequence (red), the spacer region (green) and the start codon (blue). Within the set of spacer variants, we included a variant lacking both a spacer and the core SD sequence (No RBS). Each RBS was assembled with P_V02 TSS_ and eYFP (yellow) (see **Fig. S2A**) and assessed in *E. coli* (**B**) or *Synechocystis* (**C**). For the latter, we included the P_cpc560_ promoter for comparison. Fluorescence values are shown for *E. coli* transformants at early exponential phase (5 hours) and *Synechocystis* transconjugants after 48 hours of growth, respectively. Error bars represent the ± mean SE of at least 10,000 individual cells for four biological replicates.

In *E. coli*, no eYFP fluorescence was observed in the absence of a SD sequence or a spacer region (**Fig. 3B**). However, a single nucleotide was sufficient to support translation, with peak expression observed between spacer lengths of 4-6 nt, followed by a decline with longer spacers. These results are consistent with previous reports where the optimal spacer region in *E. coli* ranged between 5-9 nt, with a maximum peak reported at 5 nt (Chen et al., 1994; Komarova et al., 2020). AU content and secondary structure within the spacer region also strongly affect translation efficiency (Allert et al., 2010). Although we did not explore those aspects here, we retained a high AU content in the spacer variants tested. We further evaluated the BioBrick-derived RBS B0032 and RBS B0033, which have reported translation efficiencies of ∼30% and ∼1% relative to RBS B0034 (iGEM Registry. Registry of Standard Biological Parts. http://parts.igem.org). Our observed eYFP fluorescence levels aligned with those characterisations, confirming the expected hierarchy of expression. The two cyanobacteria-derived RBSs, RBS cpc and RBS*, showed very poor expression in *E. coli*.

In *Synechocystis*, eYFP fluorescence across the different spacer lengths followed a trend similar to that observed in *E. coli*, with peak expression between 4–7 nt and a sharper decline with longer spacers. RBS B0034, B0032 and B0033 showed comparable expression patterns to that in *E. coli* and previous work in *Synechocystis* (Englund et al. 2016). In contrast, RBS cpc and RBS* performed markedly better than in *E. coli*. RBS cpc showed expression levels similar to RBS B0034, while RBS* produced the highest eYFP fluorescence observed. Notably, RBS* was specifically designed to be complementary to the anti-SD sequence of the *Synechocystis* 16S rRNA (Heidorn et al., 2011). Expression levels of RBS cpc were still significantly lower than the cpc560 promoter, demonstrating the strength of the upstream transcription factor binding sites in the latter sequence compare to P_V02 TSS_ (Zhou et al., 2014).

Excluding RBS cpc and RBS*, eYFP fluorescence showed a strong correlation between *E. coli* and *Synechocystis* across the RBSs tested (R^2^ = 0.95) (**Fig. S2B**). Thus, within this dataset, spacer length appeared to have a similar impact on translation in *E. coli* and *Synechocystis*. However, RBS performance is reported to vary between cyanobacteria and *E. coli* (Heidorn et al., 2011; Englund et al. 2016; Wang et al., 2018), and different combinations of promoters, RBSs and coding sequences can substantially influence translation efficiency (Salis et al., 2009; Thiel et al., 2018; Gupta and Srivastava, 2021; Jodlbauer et al., 2024). Overall, our data demonstrate that CyanOperon is well positioned as a flexible platform to systematically dissect such promoter–RBS–coding sequence interactions and enable predictive tuning of translation.

### Assembly and expression of fluorescent operons in Synechocystis

To validate the functionality of CyanOperon for operon expression in a cyanobacterium, we designed and assembled a suite of vectors to express up to three operonic fluorescent proteins in *Synechocystis*, selected to have minimal overlap in excitation/emission spectra (**Fig. 4A**). We tested expression from self-replicating RSF1010-derived vectors and following genomic integration into neutral site locus slr0397 (NS4 in Vasudevan, et al., 2019; N16 in Pinto et al., 2015). For the self-replicating plasmids, we also expressed these in *E. coli* for comparison.

**Figure 4.**
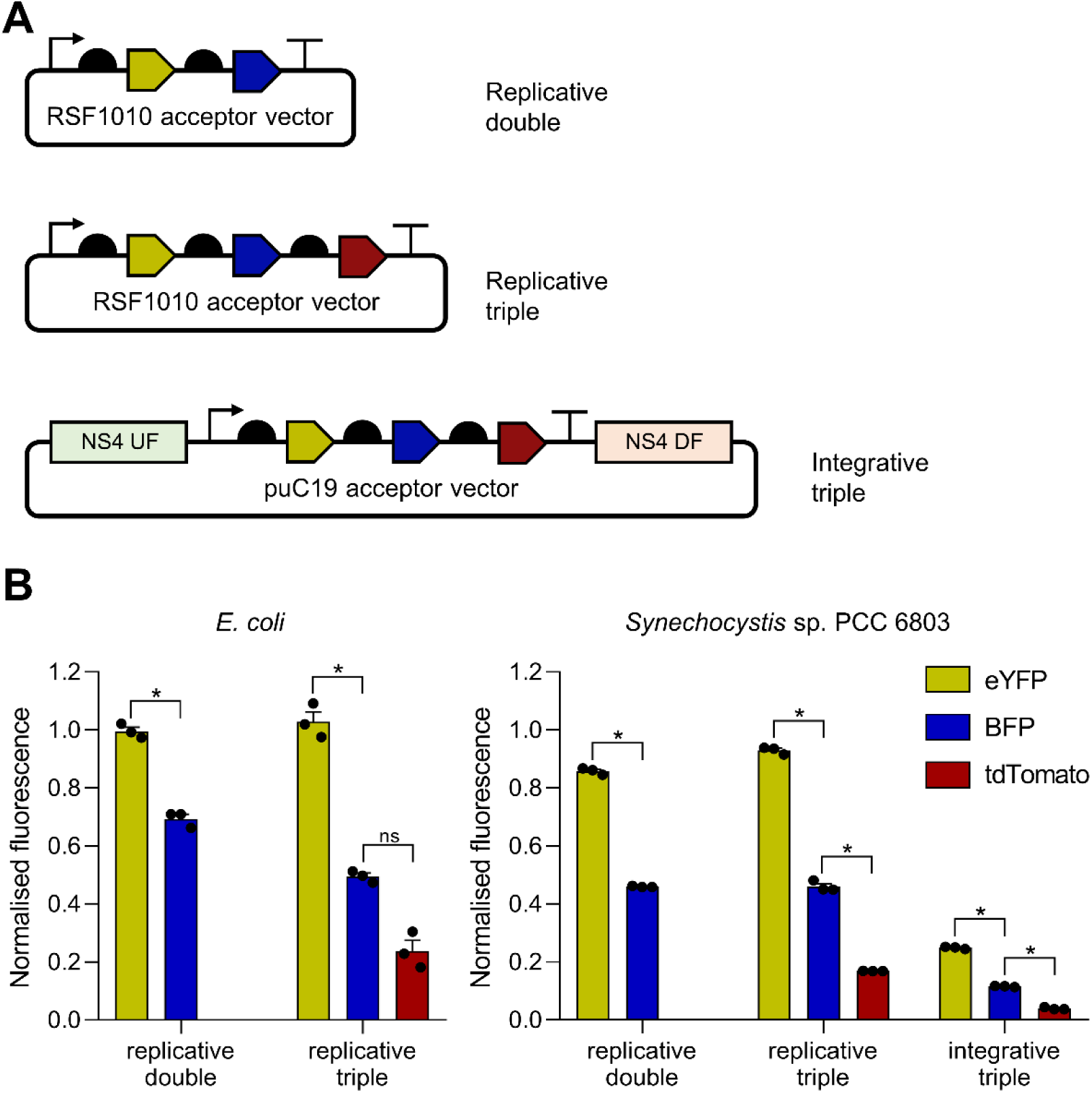
Expression of operonic fluorophores. (**A**) eYFP, BFP and tdTomato were assembled as operons into the self-replicating RSF1010-derived pPMQAK1-T vector or integrative pUC19-T vector flanked by NS4 homology arms (NSF UF and NS4 DF) for genomic integration into *Synechocystis*. (**B**). Fluorescence values for each operon-containing strain were normalised to *E. coli* or *Synechocystis* strains expressing a single fluorophore as a monocistronic unit. Normalised fluorescence values are shown for *E. coli* transformants at early exponential phase (5 hours) and *Synechocystis* transconjugants and transformants after 48 hours of growth, respectively. Error bars represent the ± mean SE of at least 100,000 individual cells for three biological replicates. Asterisks indicating significant difference (*P <* 0.05), as determined by Student’s t-test or ANOVA followed by Tukey’s honestly significant difference test. Abbreviation: ns, not significant.

In both *Synechocystis* and *E. coli*, a decline in fluorescence signal was observed for each successive fluorescent protein in both two- and three-gene operons (**Fig. 4B**). This consistent gradient likely reflects constraints inherent to the synthetic operon architecture rather than host-specific regulatory differences. Such expression dampening could arise from premature transcription termination, the accumulation of RNA degradation intermediates, or translational coupling effects (Newbury et al., 1987; Rex et al., 1994; Peters et al., 2011; Levin-Karp et al., 2013; Jeon et al., 2025). Although transcription termination and RNA degradation differ between *Synechocystis* and *E. coli*, for example, cyanobacteria lack a homologue of the *E. coli* termination factor Rho and possess distinct RNase complements (D’Heygère et al., 2013; Hui et al., 2014; Cavaiuolo et al., 2020), the similar positional expression profiles observed in both hosts suggest a broadly conserved positional effect within these synthetic operons.

In *Synechocystis*, the three-gene operon integrated at the NS4 locus displayed a similar expression pattern to that observed from the self-replicating RSF1010 vector. However, overall fluorescence was reduced by approximately 4-fold, consistent with previous findings that the RSF1010 copy number is ∼2-3-fold higher than the chromosomal copy number in *Synechocystis* during exponential growth (Nagy et al., 2021).

## Conclusions

CyanOperon expands the CyanoGate molecular cloning kit to enable modular, hierarchical assembly of multi-gene operons. We have demonstrated its utility by expressing the violacein pathway and multiple fluorescent proteins. We also compared an RBS library in *E. coli* and *Synechocystis*, revealing both conserved trends and host-specific differences in expression strength, particularly for native cyanobacterial RBSs. The observed host-dependent differences in translational performance reinforce the need for strain-specific optimisation and iterative design–build–test–learn cycles in cyanobacterial engineering. By providing a standardised framework compatible with existing MoClo systems, CyanOperon lowers the technical barrier to multi-gene operon construction and accelerates the development of engineered cyanobacterial strains for metabolic engineering and engineering biology applications.

## Methods

### Cyanobacterial culture conditions

*Synechocystis* sp. PCC 6803 (*Synechocystis*) was cultured in a standard BG-11 medium (Vasudevan et al., 2019). Transconjugant strains were cultured in BG11 supplemented with 50 µg/ml kanamycin. Liquid cultures were grown under continuous light at 30°C, 100-150 μmol photons m^-2^ s^-1^ and shaking at 120 rpm in a Multitron Pro incubator supplied with warm white LED lighting (Infors HT) or in an Algaetron AG230 (PSI). Agar plates were incubated under identical conditions, without shaking.

### Cyanobacterial conjugation and transformation

Genetic modification by conjugal transfer in *Synechocystis* was facilitated by *E. coli* strain MC1061 carrying the mobilizer vector pRK24 (www.addgene.org/51950) and helper vector pRL528 (www.addgene.org/58495) (Gale et al., 2019). Transformation with integrative Level T vectors was performed as in Vasudevan et al. (2019) using neutral site flanks pC0.171 (https://www.addgene.org/119639) and pC0.172 (https://www.addgene.org/119640).

### Vector construction and parts assembly

All cloning was performed in OneShot TOP10 *E. coli* cells. Transformed cells were cultured in LB medium and on 1.5% (w/v) LB agar plates supplemented with either 100 µg/ml spectinomycin, 100 µg/ml carbenicillin, or 50 µg/ml kanamycin as required. *E. coli* strain MC1061 was cultured in LB medium supplemented with 100 μg/ml ampicillin and 25 μg/ml chloramphenicol. All *E. coli* strains were grown at 37°C with shaking at 225 rpm.

### Construction of acceptor vectors

The Level 1 operon acceptor vectors (pOP1-prom and pOP2-6) were modified from vectors pICH47732, pICH47742, pICH47751, pICH47761, pICH47772 and pICH47781 (Engler et al., 2014). The BsaI sites of the vectors were modified to GGAG-GCTT (pOP1) or TACT-GCTT (pOP2-6) (**Fig. 1**, **Table 1**). The Level 1 acceptors pUpFlank and pDwnFlank were generated from pOP1-prom and pICH47732, modifying the BpiI sites to be compatible with CT.703 and pOP1 and any pTL and CT.703 respectively. The terminator end-linker vectors (pTL1-6) were modified from vectors pICH50872, pICH50881, pICH50892, pICH50900, pICH50914 and pICH50927 (Engler et al., 2014), with the BsaI sites modified to GCTT-CGCT. The Level T acceptor pUC19-T was generated from an available pUC19 backbone, with BpiI sites CCGC-CCAT for assembly with pUpFlank and pDwnFlank.

### Assembly of vectors for expressing violacein

Gene coding sequences for VioA, VioB, VioD and VioE were amplified from vector pMA_Level1Α_VioABDE (Andreou and Nakayama, 2018), and assembled into the Level 0 CDS1 acceptor vector pICH41308 (Engler et al., 2014). The gene coding sequence for VioC was synthesised (Integrated DNA Technologies) and cloned into pICH41308.

For Level 1 assembly, VioA was assembled in the pOP1 acceptor vector with TSS promoter P_J23100_, P_J23101_, P_J23103_, P_J23105_, P_J23110_ or P_V02_ (Vasudevan et al., 2019), and RBS B0034. VioB, VioD, VioE and VioC were assembled with RBS B0034 in acceptor vectors pOP2, pOP3, pOP4 and pOP5, respectively. The T_rrnB_ double terminator was assembled into the pTL5 acceptor vector. The Level 1 vectors were then assembled into the Level T acceptor pPMQAK1-T (pCAT.000).

### Construction of TSS promoter and RBS vectors

TSS promoters were based on existing promoters in the CyanoGate library (Vasudevan et al., 2019) and constructed by annealing complementary oligonucleotides (Sigma Aldrich) for assembly into the Level 0 TSS acceptor pCA0.324 (**Table 1**; **Table S1**). The sequences were flanked with BpiI restriction sites generating overhangs GGAG and TACT, different to those of the TSS acceptor in the CyanoGate library (GGAG-TAGC, pCA0.001). Hence, CyanOperon TSS promoter parts are labelled V2 (**Materials S1**). All RBS sequences were assembled into the Level 0 TSS acceptor pCA0.325 by annealing complementary oligonucleotides as above (**Table S1**). The sequences were flanked with BpiI restriction sites generating overhangs TACT and AATG (**Fig. 3A**). Level 1 poP1-prom vectors were assembled with P_V02 TSS_ (pC0.335), eYFP (pC0.008) and each RBS part. The T_rrnB_ double terminator was assembled into the pTL1 acceptor vector. Level 1 vectors were assembled into pPMQAK1-T.

### Assembly of Level 1 and Level T plasmids for fluorescent operon constructs

Level 0 vectors carrying fluorescent proteins eYFP, BFP or tdTomato were assembled with TSS promoter P_V02 TSS_ and RBS B0034 in pOP1-prom (eYFP), or with RBS B0034 in pOP2 (BFP) or pOP3 (tdTomato). Level 1 flank vectors for genome integration at the NS4 locus were constructed by assembling 6803 NS4 up flank (pC0.171) and the unmark linker (pC0.116) into pUpFlank and 6803 NS4 down flank (pC0.172), AbR up linker (pC0.121) and the spectinomycin resistance cassette (pC0.028) into pDwnFlank. The T_rrnB_ double terminator was assembled into the pTL1-3 acceptor vectors. Level 1 vectors were assembled into the Level T acceptor pPMQAK1-T or pUC19-T.

### Violacein extraction and quantification

Transgenic *E. coli* seed cultures were grown from single colonies in LB medium with appropriate antibiotics at 37 °C with shaking at 250 rpm for 12 hr. Seed cultures were then inoculated into 20 ml fresh LB medium and grown at 37 °C for 24 hr. Cultures (2 ml) were pelleted at 17,000 x g for 10 min. Violacein was then extracted in 100% (v/v) ethanol at 95 °C in a heat block for 5 min. The extract was pelleted at 17,000 x g for 10 min to remove cell debris. Violacein concentration was determined spectrophotometrically at 575 nm using the extinction coefficient of 56.01 mL mg^-1^ cm^-1^ for pure violacein in 100% (v/v) ethanol (Mendes et al., 2001).

### Fluorescence assays

Transgenic *E. coli* seed cultures were diluted 1:1000 in flat-bottom black 96-well plates (Greiner BioOne) and grown for 5 hr at 37 °C with shaking at 250 rpm. The optical density at 600 nm (OD_600_) was measured using a FLUOstar OMEGA micro-plate reader (BMG Labtech).

Seed cultures of transconjugated *Synechocystis* strains were grown in BG11 supplemented with 50 µg/ml kanamycin to an OD_750_ = 1-2. The seed cultures were diluted to OD_750_ = 0.2 in 24-well clear plates (Costar Corning Incorporated) and grown in a Multitron Pro incubator as described above.

For RBS characterisation, eYFP fluorescence for *E. coli* or *Synechocystis* samples (10,000 cells per culture) were measured by flow cytometry using a FACSCanto II Flow Cytometer (Becton Dickinson). For the fluorescent operons, fluorescence for eYFP, mBFP and tdTomato for individual *E. coli* or *Synechocystis* cells (100,000 cells per culture) was measured by flow cytometry using an LSRFortessa Flow Cytometer (Becton Dickinson). Cells were gated using forward and side scatter, and median eYFP, mBFP or tdTomato fluorescence was calculated from excitation/emission wavelengths 488 nm/525 nm, 405 nm/450 nm or 550 nm/580 nm, respectively (Vasudevan et al., 2019).

## Acknowledgments

The authors thank Chris Hall and Marie Geopp for their technical assistance with flow cytometry.

## Supplementary Tables and Figures

**Table S1.**
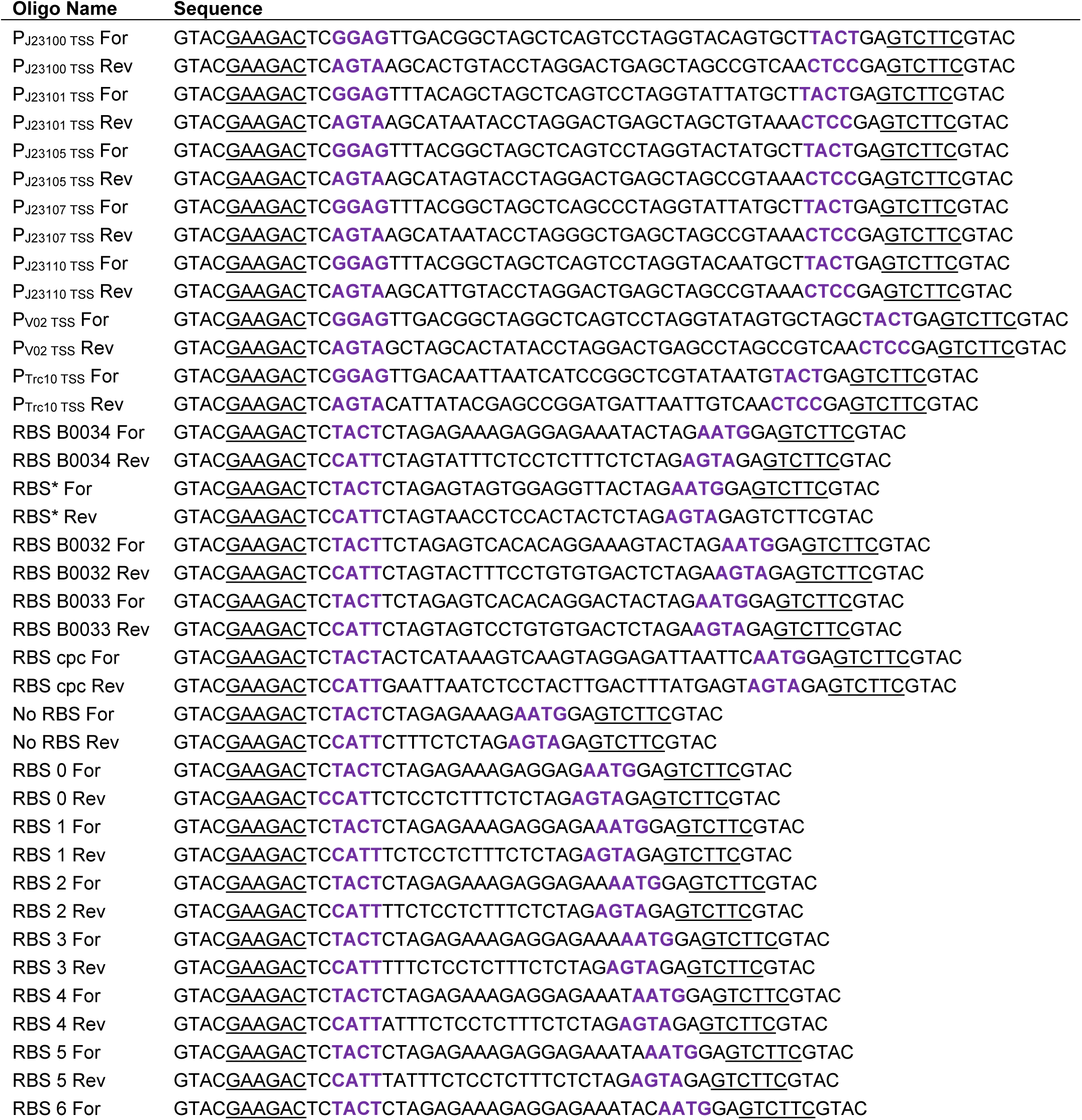
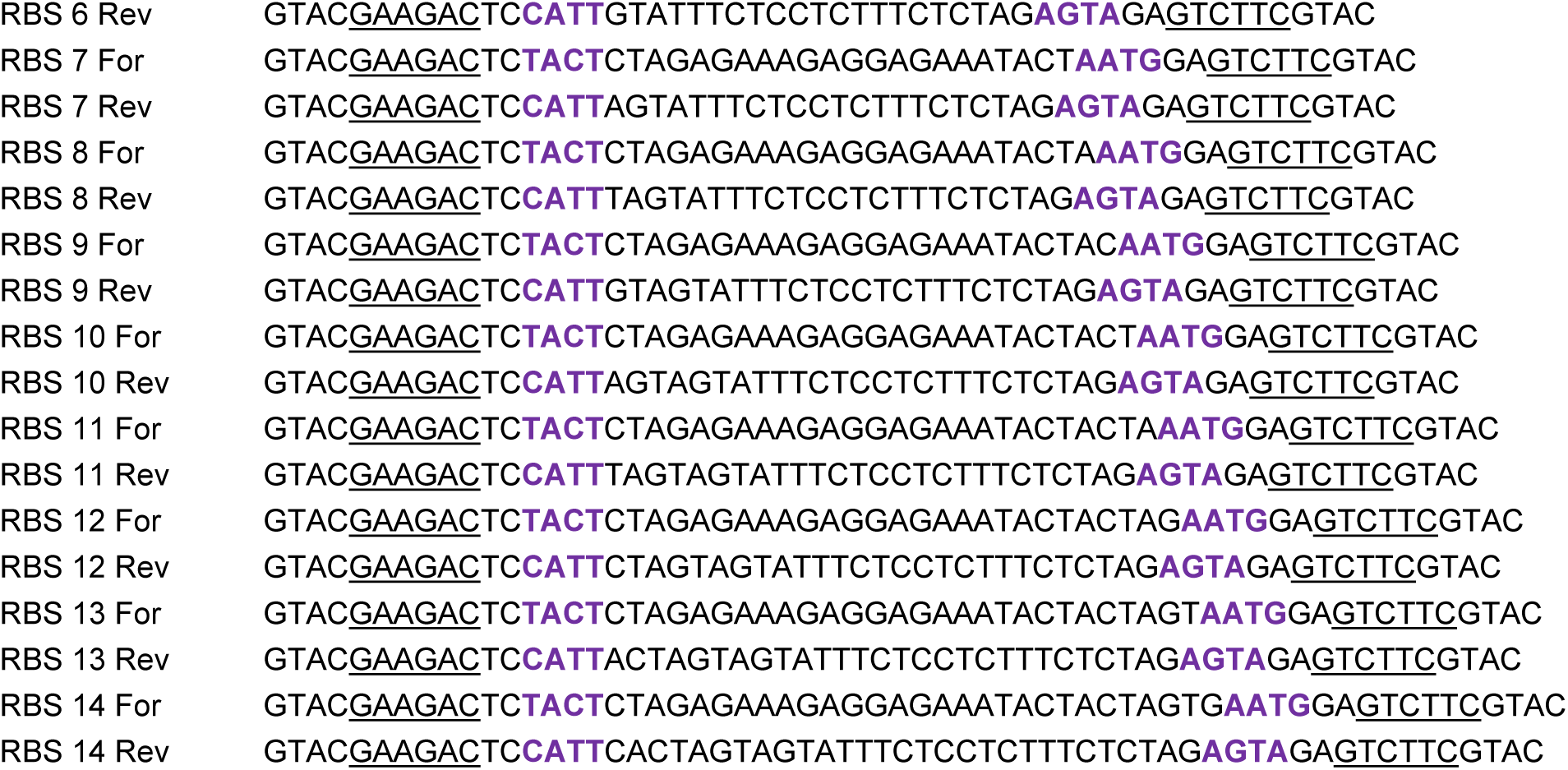
Oligonucleotides used to assembly Level 0 TSS promoter and RBS parts. Complementary oligonucleotides were annealed by mixing equal volumes in equimolar amounts, incubating the mixture at 95°C for 5 minutes, and cooling slowly to room temperature. The annealed oligo mixture was used directly in MoClo assembly reactions (Vasudevan et al. 2019).

**Figure S1.**
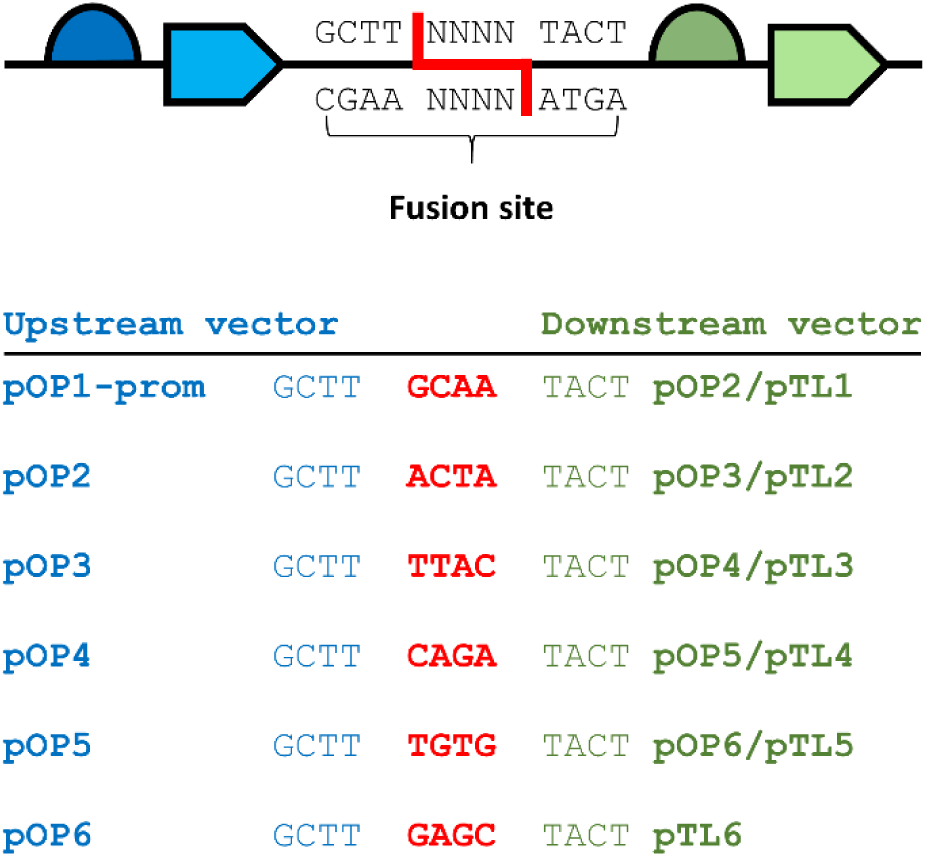
Overview of the fusion sites between assembled pOP vectors. The 12 bp intergenic regions between the coding sequence and the subsequent next ribosome binding site in the operon are shown for pOP1-prom to pOP6 (or the appropriate Terminator end-linker (pTL) vector). Following Level T assembly the fusion site consists of two legacy 4 bp scars from previous Level 1 vector assemblies with BsaI (GCTT and TACT in blue and green, respectively) and a unique 4 bp scar depending on the level positions being assembled (shown in red).

**Figure S2.**
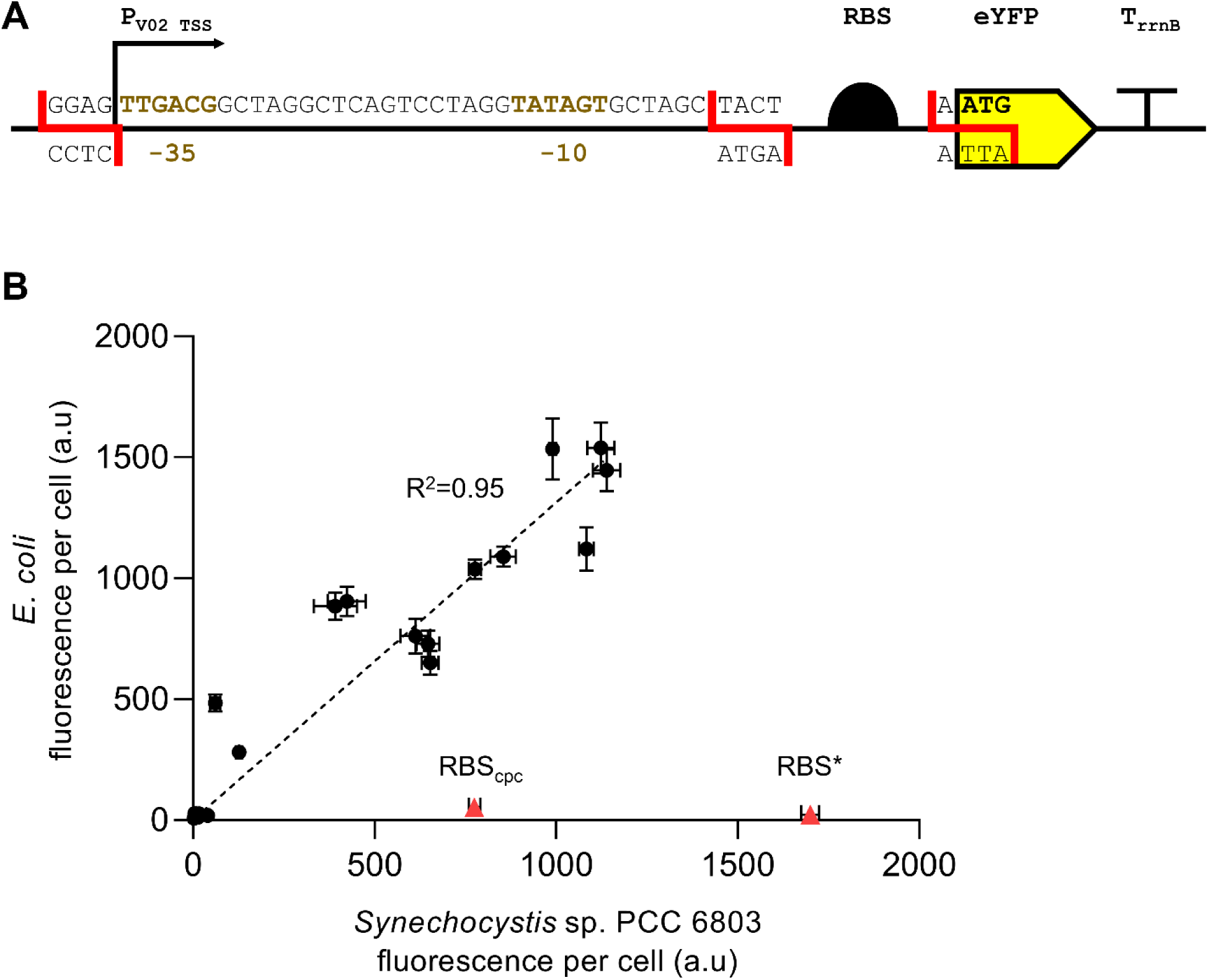
Format of the expression cassettes assembled to assess RBS strength and correlation analysis. **A.** The sequence of the P_V02 TSS_ promoter (pC0.335), is shown with the -35 and -10 elements highlighted. The common 4 bp overhangs used for assembly are shown in red. **B.** Correlation analysis of fluorescence data for *E. coli* and *Synechocystis* sp. PCC 6803 from Fig. 3. RBS cpc and RBS* (red) were considered outliers and excluded from the coefficient of determination (R^2^) analysis.

**Materials S1. Plasmid maps of vectors in Table 1**. All plasmids are available on Addgene.

**Materials S2. Plasmids maps of assembled RSF1010-derived vectors (pPMQAK1) carrying the VioABCDE operon.** Expression is driven by six different promoters and RBS B0034 (P*_x_* _TSS_–RBS B0034–VioA– RBS B0034–VioB–RBS B0034–VioE–RBS B0034–VioD–RBS B0034–VioC–T_rrnB_).

